# Coccinellid host morphology dictates morphological diversity of the parasitoid wasp *Dinocampus coccinellae*

**DOI:** 10.1101/460998

**Authors:** Hannah Vansant, Yumary M Vasquez, John J Obrycki, Arun Sethuraman

**Affiliations:** Department of Biological Sciences, California State University San Marcos, San Marcos CA 92096; Department of Entomology, University of Kentucky, Lexington KY 40546; These authors contributed equally to this work; Current address: Graduate Program in Quantitative and Systems Biology, University of California Merced, Merced CA 95343; Current address: NIH MHIRT Program, University of Oxford Oxford, OX1 3PT

**Keywords:** Morphology, parasitoid-host interaction, phenotypic plasticity

## Abstract

Pararsitoid-host interactions involving host species that are newly introduced into the range of a generalist parasitoid provide systems that can be examined for phenotypic plasticity and evolutionary changes in parasitoid-host dynamics. The solitary Braconid parasitoid wasp, *Dinocampus coccinellae*, has a cosmopolitan distribution and parasitizes approximately 50 species of predatory lady beetles (ladybirds) in the family Coccinellidae. In this study we quantified the effect of six (4 native North American and 2 non-native North American) host species on the morphometrics of *D. coccinellae*. Adult lady beetles were collected from 13 locations in the United States and reared in the laboratory until *D.coccinellae* exited from their adult beetle hosts. Eighty-nine individual *D. coccinellae* females and their associated host were weighed and morphometric measurements were taken. The smallest lady beetle host *Hippodamia parenthesis* produced the smallest adult wasps; the largest host species, *Coccinella septempunctata*, produced the largest female wasps. A directional cline in morphology of wasps and their coccinellid hosts was also observed in a dry-weight regression (R^2^ = 0.4066, p-value < 0.0001). Two underlying mechanisms may explain the results of our study: (1) morphometric variation in *D. coccinellae* is governed by phenotypic plasticity with the size of the emerging offspring contingent on the size of the coccinellid host, and/or (2) that morphometric variation in *D. coccinellae* is governed by genomic adaptation to coccinellid host populations.

## Introduction

Parasitic Hymenoptera make up at least 280,000 species of all parasitic insects (Pennacchio and Strand 2006). Numerous studies have examined the relationship between hosts and their parasitoids. (Hochberg and Ives 2000, Godfray 1994). Importantly, several studies have shown that the morphological characteristics of adults and fecundity of female parasitoids are affected by host characteristics, for example, host species (Nicol et al., 1999), host life stage (Traynor et al., 2005), host instars (Cloutier et al., 2000), and host size (Harvey et al., 2006; Mackauer et al., 2001). The size and species of hosts have been shown to greatly influence the evolutionary history of morphological characteristics in parasitoids (Brandl et al., 1987; Belshaw et al., 2003; Charnov et al., 1984; Bakker K et al., 1985; Moore et al., 2002; Symonds et al., 2013). It has been hypothesized that host-parasitoid co-evolution could eventually lead to increased fitness of parasitoids allowing them to parasitize multiple species (Charnov et al.,1984; Ellers et al., 2002; Sampaio et al., 2008). This also points to evidence of host-specificity in a majority of parasitoids, in that parasitoids adaptively evolve in response to host characteristics, and eventually may specialize in parasitizing only particular host species. Few studies examine generalist parasitoids that parasitize multiple host species, and correspondingly exhibit quantitative variability in morphology, potentially in response to host characteristics. For example, parasitoid size as a plastic trait has been studied using *Aphidus ervi (Hymenoptera:Aphidiidae)*, an aphid parasitoid, that attacks multiple host species (Henry et al., 2006). There is little understanding, however, of the biological processes that lead to generalist behavior of parasitoids.

*Dinocampus coccinellae* (Hymenoptera: Braconidae) is a thelytokous parthenogenic species, in which females are produced from unfertilized eggs (Ceryniger et al 2012). Males have been rarely observed; one laboratory study observed mating to occur, but all offspring were females (Wright 1980). This process results in offspring being maternal clones. *D. coccinellae* are generalist wasps, capable of parasitizing over 50 species of coccinellids across various climates worldwide (Balduf 1926, Ceryniger et al 2012). Within the beetle host, *D. coccinellae* larvae feed on teratocytes derived from the parasitoid egg, thus the adult beetle host typically survives the larval development of *D. coccinellae* (Ceryniger et al 2012). However, most parasitized adult beetles die following the exit of the parasitoid larva, when they become entangled in the pupal coccoon produced by the parasitoid (Ceryniger et al 2012, Dheilly et al 2015). Several predatory lady beetle hosts of *D. coccinellae* are natural enemies that are beneficial species for biological control (including the native North American species *Hippodamia convergens*, and two non-native species in North America, *Coccinella septempunctata*, and *Harmonia axyridis)*. Coupled with low survival rates of parasitized beetles, and the generalist nature of *D. coccinellae*, these wasps are of biological, ecological, and economic interest. Specifically, we are interested in examining the ecological basis of host-specific plastic or adaptive morphological traits, that make *D. coccinellae* amenable to parasitizing coccinellid beetles. In this study we analyze the geometric morphometrics of field-collected, lab-reared *D. coccinellae* and their coccinellid beetle hosts (from six host species), sampled across the United States. Our goals in this study were twofold: (1) to quantify the variability in morphometrics of *D. coccinellae* across its primary range in the United States, (2) to correlate the variability in morphometrics of *D. coccinellae* with morphometrics of their hosts. Broadly, we hypothesize that morphometric diversity of the host species will dictate the morphometrics of the parasitoid wasps parasitizing them.

## Methods

Ninety-nine parasitoid wasps within their adult hosts (*Harmonia axyridis* (*Har. axyridis*), *Coleomegilla maculata* (*Col. maculata*), *Coccinella septempunctata* (*Cocc. septempunctata*), *Hippodamia convergens* (*H. convergens*), *Cycloneda munda* (*Cyc. munda*), and *Hippodamia parenthesis* (*H. parenthesis*)) were field-collected in the states of Kentucky, Ohio, Illinois, New York, Missouri and Kansas (see Fig. 1) The Hippodamia convergens samples from Arizona were field-collected and shipped to the authors. Adult lady beetles were collected from agricultural fields, prairie, and roadside vegetation using sweep nets. Predatory Coccinellidae are commonly found in these habitats when their aphid prey is present. Following field collection, adult beetles were reared in the laboratory (L:D 16:8, 22°C, on pea aphids) until the last larval stage of the wasps’ development at which point the parasitoid larva exits the host and pupates within a cocoon, typically woven between the coccinellid beetle hosts’ legs. After eclosion of the adult parasitoid, both the beetle host and parasitoid were then stored in 95% ethanol at −20C. Of these samples, 89 were viable for the morphological analyses which needed intact, undamaged samples of both the parasitoid and host. It is of interest to note that of the six host species of coccinellids, *Harmonia axyridis* and *Coccinella septempunctata* are not native to continental United States (derived from Asia, and Europe respectively, (Obrycki and Kring 1988), thus the interaction between these host species and North American *D. coccinellae* may be a recently evolving interaction. Alternately, *D. coccinellae* could also be native to Europe or Asia (Ceryniger et al 2012), and have shifted hosts since their introduction to North America. Dry weights of wasps and host were recorded individually on a Mettler Toledo XS105 DualRange Analytical Balance after 1 minute of air-drying on a Kimtech Kimwipe to allow for evaporation of alcohol. After weighing, each wasp and its respective host were photographed individually in two replicate rounds using an optical microscope with a SPOT Idea camera attachment. Wasps were photographed from a lateral view (see Fig. 3), and their hosts were photographed from lateral, dorsal, and ventral views to include key morphological characteristics and maintain consistency in imaging (see Fig. 2). Images were uploaded into Image-J (version 1.51j8) for morphometric measurement (in mm) of host and wasp morphological characteristics with a scale bar (included in the parasitoid/host mounting stage) to ensure consistent scaling. Wing length of the wasp was not included in further statistical analyses due to many wings being folded, or crushed during storage. Body depth of hosts (measured as the “height” of a beetle from the lateral view) was also excluded due to measurement inconsistency on the styrofoam mounting stage.. All high-resolution images from this study will be deposited with http://www.morphbank.net/ upon acceptance.

**Figure 1.**
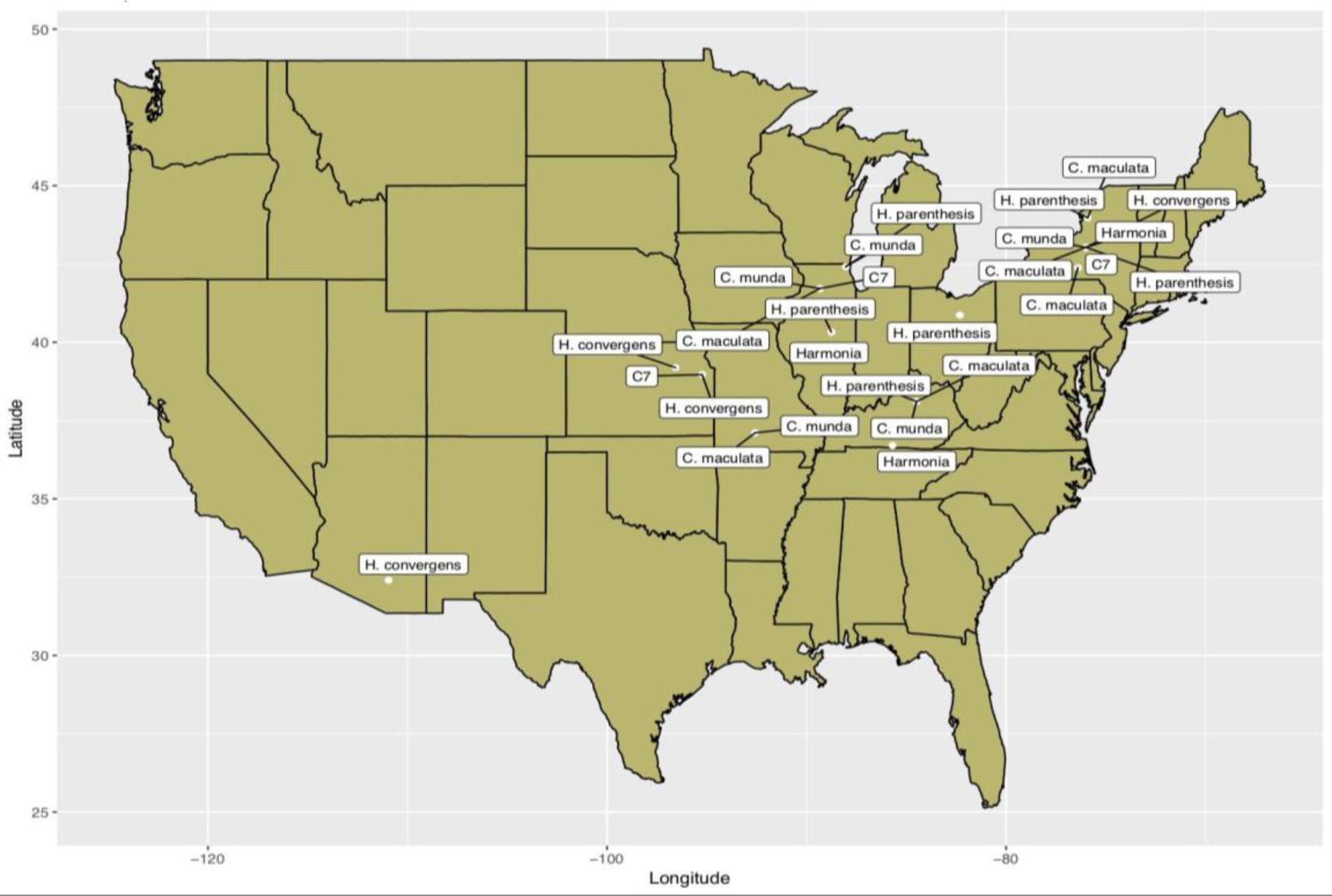
Map of locations from which Coccinellid hosts with their parasitoid wasp were field-collected (except *H. convergens* from Arizona, which was shipped to the authors).

**Figure 2.**
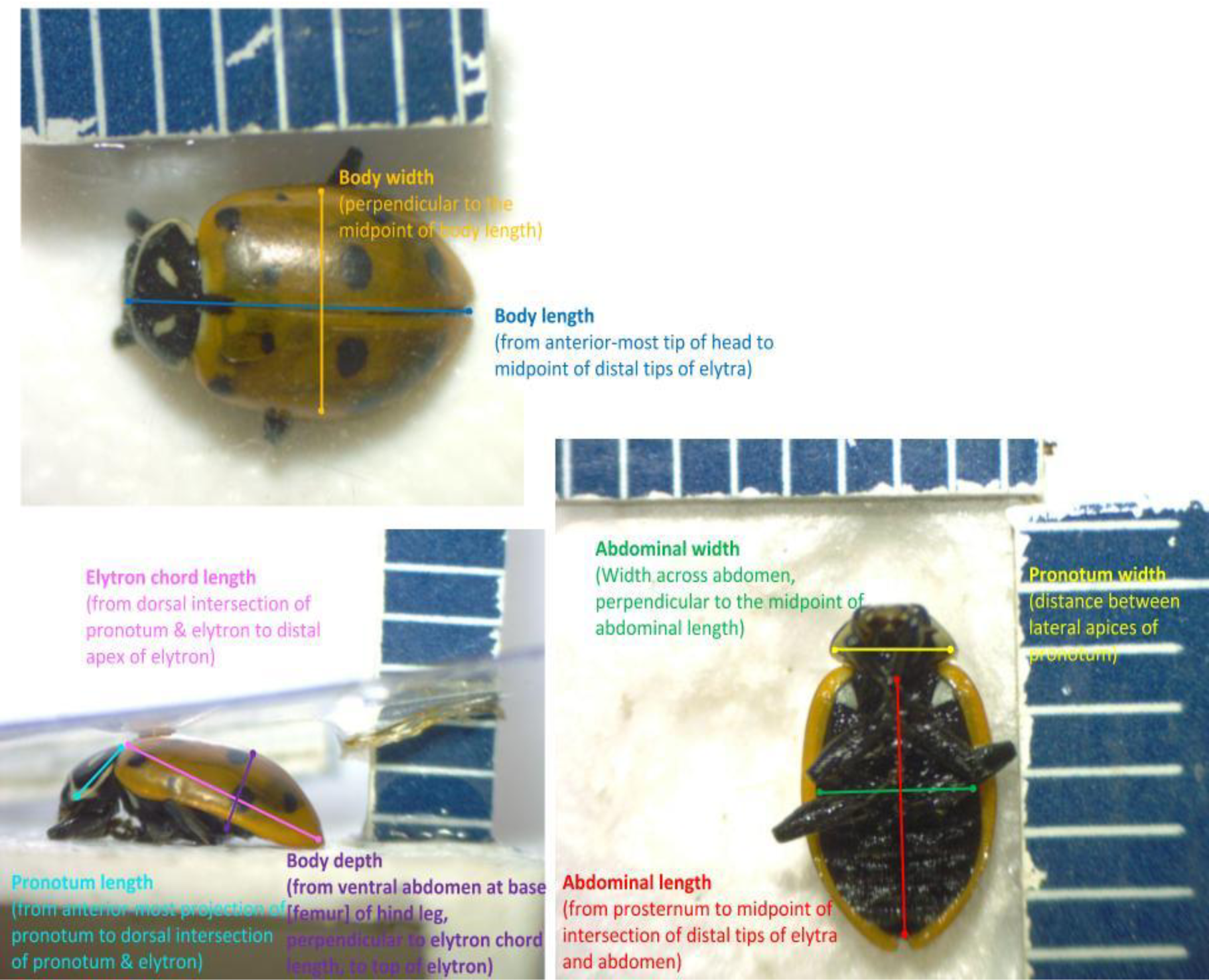
Morphometric variables measured from Coccinellid hosts, shown in three perspectives - dorsal, lateral, and venral views. Image courtesy: D. Sustaita.

**Figure 3.**
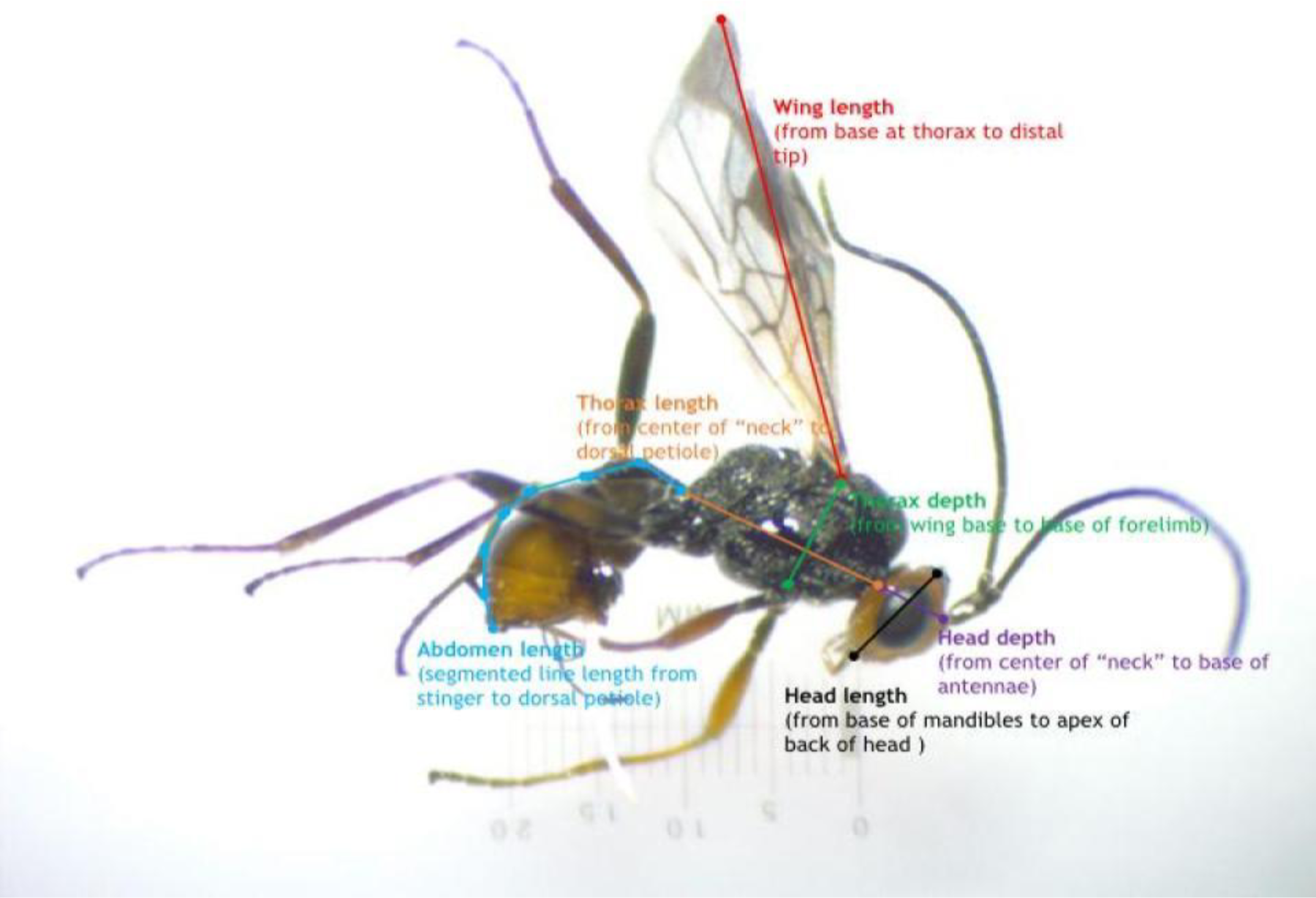
Morphometric variables measured from parasitoid wasps, *D. coccinellae*, shown in a single lateral perspective. Image courtesy: D. sustaita.

All statistical analyses of morphometric measurements were carried out for each round separately (except for dry weight which was recorded in one round) using Rstudio (version 1.0.143) and the package: ggplot2. Variation in means of recorded morphometrics of wasps and their hosts were visualized using boxplots, and summaries (means, medians, standard deviations) were computed. Due to lack of normality across most morphometric measures, a Wilcoxon signed rank test was performed across wasp morphometric measures (grouped by their respective host species) to test the null hypothesis of non-significant deviation from the mean. Similar Wilcoxon signed rank tests were also performed on coccinellid host morphometrics. We then performed a Kruskal-Wallis non-parametric one way ANOVA, followed by a posthoc Dunn’s Test on each morphometric measure in the parasitoid versus its host species, to test for significant morphometric differences by host. Parasitoid dry-weight was also regressed onto host dry-weight, as a proxy of size variation between the two. Additionally, a Principal Components Analysis (PCA) was used to orthogonally transform morphometrics (including dry-weights) of parasitoids and their hosts, and project their variability onto principal components of maximum variance.

## Results

Summaries of morphometric measurements in coccinellid hosts (Table 1), and parasitoid wasps (Table 2) are shown in Figures 4 and 5. The smallest hosts (Fig. 4 as judged by all measurements), *H. parenthesis* produced the smallest adult wasps (Fig. 5). Similarly, the largest hosts, *Coccinella septempunctata*, produced the largest wasps. This directional cline (Fig. 6) in morphology of wasps and their coccinellid hosts is also reflected in a dry-weight regression (R^2^ = 0.4066, p-value < 0.0001). A non-parametric Kruskal-Wallis one-way analysis of variance (ANOVA) test of host morphometric variation and parasitoid wasp morphometric variation was significant at a p-value threshold of 0.05 on each morphometric measurement, rejecting the null hypothesis that there is no variation in means of measured morphometric variables in parasitoid wasps, among their corresponding host species. Additionally, coccinellid host measurements were significantly variable (p < 0.001) among all host species measured, indicating significant variability in sizes of coccinellid hosts. Conservative posthoc Dunn’s tests indicated that the variability observed between pronotum width in coccinellid adult hosts had the greatest level of significance. A similar post-hoc Dunn’s test in parasitoid wasp measurements indicated that thorax length varied more than other morphometric measurements. A Principal Components Analysis (PCA) showed that the first two PC’s account for 70-75% of variability in both rounds (Fig. 7). Interestingly, morphometric variables in the two introduced beetle species, *Cocc. septempunctata* and *Har. axyridis* share no overlapping points with the native species, *Col. maculata*, *Cyc. munda*, and *H. parenthesis*, but show close association with the native species *H. convergens.*

**Table 1.** Summary of Coccinellid host (*H. parenthesis, H. convergens, C. maculata, C. septempunctata, H. axyridis, C. munda*) morphometric measurements from the first round of measurements, including abdominal widths, abdominal lengths, pronotum widths, elytron chord lengths, pronotum lengths, body lengths, and body widths, in millimeters. The p-values are derived from Wilcoxon signed-rank tests for significant deviation from mean morphometric measurements within each host species class.

|  | Host Length |  |  |  |  |  |  |
| --- | --- | --- | --- | --- | --- | --- | --- |
| Coccinellid Host |  |  |  |  |  |  |  |
|  | Abdominal Width | Abdominal Length | Pronotum Width | Elytron Chord Length | Pronotum Length | Body Length | Body Width |
| *H. parenthesis* | 2.47 (sd = 0.2, p-value = 2.384e-07, min = 1.92, median = 2.48, max = 2.76,) | 3.42 (sd = 0.22, p-value = 2.886e-05, min = 2.82, median = 3.43, max = 3.7,) | 1.98 (sd = 0.15, p-value = 2.883e-05, min = 1.64, median = 2.01, max = 2.24,) | 3.85 (sd = 0.25, p-value = 2.384e-07, min = 3.21, median = 3.96, max = 4.22,) | 1.24 (sd = 0.1, p-value = 2.886e-05, min = 1.05, median = 1.23, max = 1.47,) | 4.9 (sd = 0.39, p-value = 2.384e-07, min = 4.26, median = 4.81, max = 5.76,) | 2.91 (sd = 0.2, p-value = 2.384e-07, min = 2.4, median = 2.93, max = 3.24,) |
| *H. convergens* | 3.18 (sd = 0.27, p-value = 0.001656, min = 2.77, median = 3.06, max = 3.7,) | 4.41 (sd = 0.42, p-value = 0.0002441, min = 3.84, median = 4.35, max = 5.22,) | 2.41 (sd = 0.18, p-value = 0.0002441, min = 2.19, median = 2.35, max = 2.77,) | 4.81 (sd = 0.6, p-value = 0.0002441, min = 3.39, median = 4.65, max = 5.68,) | 1.54 (sd = 0.11, p-value = 0.0002441, min = 1.38, median = 1.55, max = 1.76,) | 5.96 (sd = 0.46, p-value = 0.0002441, min = 5.3, median = 5.9, max = 6.82,) | 3.78 (sd = 0.32, p-value = 0.0002441, min = 3.43, median = 3.64, max = 4.52,) |
| *C. maculata* | 2.6 (sd = 0.19, p-value = 2.328e-10, min = 2, median = 2.6, max = 2.94,) | 3.96 (sd = 0.28, p-value = 2.328e-10, min = 3.02, median = 4.01, max = 4.38,) | 2.1 (sd = 0.12, p-value = 5.63e-07, min = 1.72, median = 2.1, max = 2.37,) | 4.35 (sd = 0.35, p-value = 2.328e-10, min = 3.32, median = 4.31, max = 5.37,) | 1.3 (sd = 0.12, p-value = 5.642e-07, min = 1.06, median = 1.31, max = 1.51,) | 5.61 (sd = 0.41, p-value = 2.328e-10, min = 4.37, median = 5.72, max = 6.26,) | 3.21 (sd = 0.22, p-value = 2.328e-10, min = 2.53, median = 3.22, max = 3.66,) |
| *H. axyridis* | 4.32 (sd = 0.28, p-value = 0.125, min = 3.97, median = 4.34, max = 4.63,) | 5.21 (sd = 0.2, p-value = 0.125, min = 4.96, median = 5.24, max = 5.39,) | 3.09 (sd = 0.2, p-value = 0.125, min = 2.85, median = 3.08, max = 3.34,) | 5.79 (sd = 0.27, p-value = 0.125, min = 5.57, median = 5.73, max = 6.15,) | 1.9 (sd = 0.06, p-value = 0.125, min = 1.85, median = 1.89, max = 1.98,) | 6.81 (sd = 0.21, p-value = 0.125, min = 6.59, median = 6.79, max = 7.07,) | 5.44 (sd = 0.27, p-value = 0.125, min = 5.05, median = 5.52, max = 5.66,) |
| *C. munda* | 2.73 (sd = 0.27, p-value = 0.003906, min = 2.34, median = 2.71, max = 3.12,) | 3.36 (sd = 0.38, p-value = 0.003906, min = 2.91, median = 3.17, max = 4.00,) | 2.26 (sd = 0.18, p-value = 0.003906, min = 2.05, median = 2.23, max = 2.51,) | 4.08 (sd = 0.64, p-value = 0.003906, min = 3.38, median = 3.96, max = 5.5,) | 1.34 (sd = 0.07, p-value = 0.003906, min = 1.22, median = 1.37, max = 1.41,) | 4.58 (sd = 0.45, p-value = 0.003906, min = 3.98, median = 4.52, max = 5.28,) | 3.69 (sd = 0.3, p-value = 0.003906, min = 3.18, median = 3.69, max = 4.14,) |
| *C7* | 4.08 (sd = 0.41, p-value = 0.02225, min = 3.93, median = 3.74, max = 4.86,) | 5.19 (sd = 0.53, p-value = 0.01563, min = 4.72, median = 4.91, max = 6.17,) | 3.26 (sd = 0.34, p-value = 0.01563, min = 2.93, median = 3.07, max = 3.78,) | 5.57 (sd = 0.92, p-value = 0.01563, min = 4.21, median = 5.8, max = 6.89,) | 2.01 (sd = 0.42, p-value = 0.01563, min = 1.13, median = 2.09, max = 2.48,) | 6.85 (sd = 0.64, p-value = 0.01563, min = 6.33, median = 6.52, max = 8.1,) | 5.4 (sd = 0.44, p-value = 0.01563, min = 4.81, median = 5.54, max = 5.98,) |

**Table 2.**
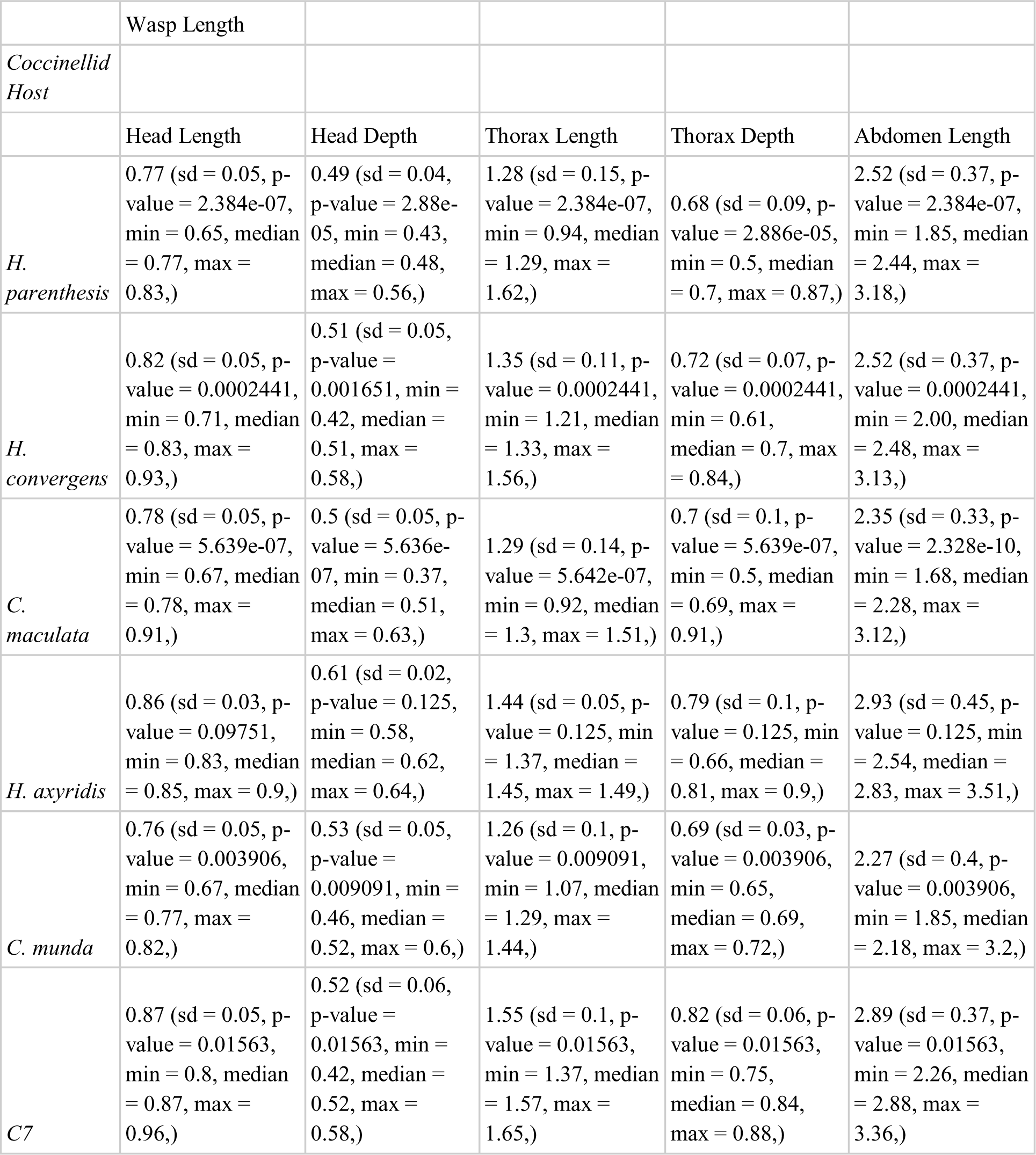
Summary of parasitoid wasp (*D. coccinellae*) morphometric measurements, including head length, head depth, thorax length, thorax depth, and abdomen length, in millimeters, from the first round of measurements. The p-values are derived from Wilcoxon signed-rank tests for significant deviation from mean morphometric measurements within each host species class.

|  | Wasp Length |  |  |  |  |
| --- | --- | --- | --- | --- | --- |
| *Coccinellid Host* |  |  |  |  |  |
|  | Head Length | Head Depth | Thorax Length | Thorax Depth | Abdomen Length |
| *H. parenthesis* | 0.77 (sd = 0.05, p-value = 2.384e-07, min = 0.65, median = 0.77, max = 0.83,) | 0.49 (sd = 0.04, p-value = 2.88e-05, min = 0.43, median = 0.48, max = 0.56,) | 1.28 (sd = 0.15, p-value = 2.384e-07, min = 0.94, median = 1.29, max = 1.62,) | 0.68 (sd = 0.09, p-value = 2.886e-05, min = 0.5, median = 0.7, max = 0.87,) | 2.52 (sd = 0.37, p-value = 2.384e-07, min = 1.85, median = 2.44, max = 3.18,) |
| *H. convergens* | 0.82 (sd = 0.05, p-value = 0.0002441, min = 0.71, median = 0.83, max = 0.93,) | 0.51 (sd = 0.05, p-value = 0.001651, min = 0.42, median = 0.51, max = 0.58,) | 1.35 (sd = 0.11, p-value = 0.0002441, min = 1.21, median = 1.33, max = 1.56,) | 0.72 (sd = 0.07, p-value = 0.0002441, min = 0.61, median = 0.7, max = 0.84,) | 2.52 (sd = 0.37, p-value = 0.0002441, min = 2.00, median = 2.48, max = 3.13,) |
| *C. maculata* | 0.78 (sd = 0.05, p-value = 5.639e-07, min = 0.67, median = 0.78, max = 0.91,) | 0.5 (sd = 0.05, p-value = 5.636e-07, min = 0.37, median = 0.51, max = 0.63,) | 1.29 (sd = 0.14, p-value = 5.642e-07, min = 0.92, median = 1.3, max = 1.51,) | 0.7 (sd = 0.1, p-value = 5.639e-07, min = 0.5, median = 0.69, max = 0.91,) | 2.35 (sd = 0.33, p-value = 2.328e-10, min = 1.68, median = 2.28, max = 3.12,) |
| *H. axyridis* | 0.86 (sd = 0.03, p-value = 0.09751, min = 0.83, median = 0.85, max = 0.9,) | 0.61 (sd = 0.02, p-value = 0.125, min = 0.58, median = 0.62, max = 0.64,) | 1.44 (sd = 0.05, p-value = 0.125, min = 1.37, median = 1.45, max = 1.49,) | 0.79 (sd = 0.1, p-value = 0.125, min = 0.66, median = 0.81, max = 0.9,) | 2.93 (sd = 0.45, p-value = 0.125, min = 2.54, median = 2.83, max = 3.51,) |
| *C. munda* | 0.76 (sd = 0.05, p-value = 0.003906, min = 0.67, median = 0.77, max = 0.82,) | 0.53 (sd = 0.05, p-value = 0.009091, min = 0.46, median = 0.52, max = 0.6,) | 1.26 (sd = 0.1, p-value = 0.009091, min = 1.07, median = 1.29, max = 1.44,) | 0.69 (sd = 0.03, p-value = 0.003906, min = 0.65, median = 0.69, max = 0.72,) | 2.27 (sd = 0.4, p-value = 0.003906, min = 1.85, median = 2.18, max = 3.2,) |
| *C7* | 0.87 (sd = 0.05, p-value = 0.01563, min = 0.8, median = 0.87, max = 0.96,) | 0.52 (sd = 0.06, p-value = 0.01563, min = 0.42, median = 0.52, max = 0.58,) | 1.55 (sd = 0.1, p-value = 0.01563, min = 1.37, median = 1.57, max = 1.65,) | 0.82 (sd = 0.06, p-value = 0.01563, min = 0.75, median = 0.84, max = 0.88,) | 2.89 (sd = 0.37, p-value = 0.01563, min = 2.26, median = 2.88, max = 3.36,) |

**Figure 4.**
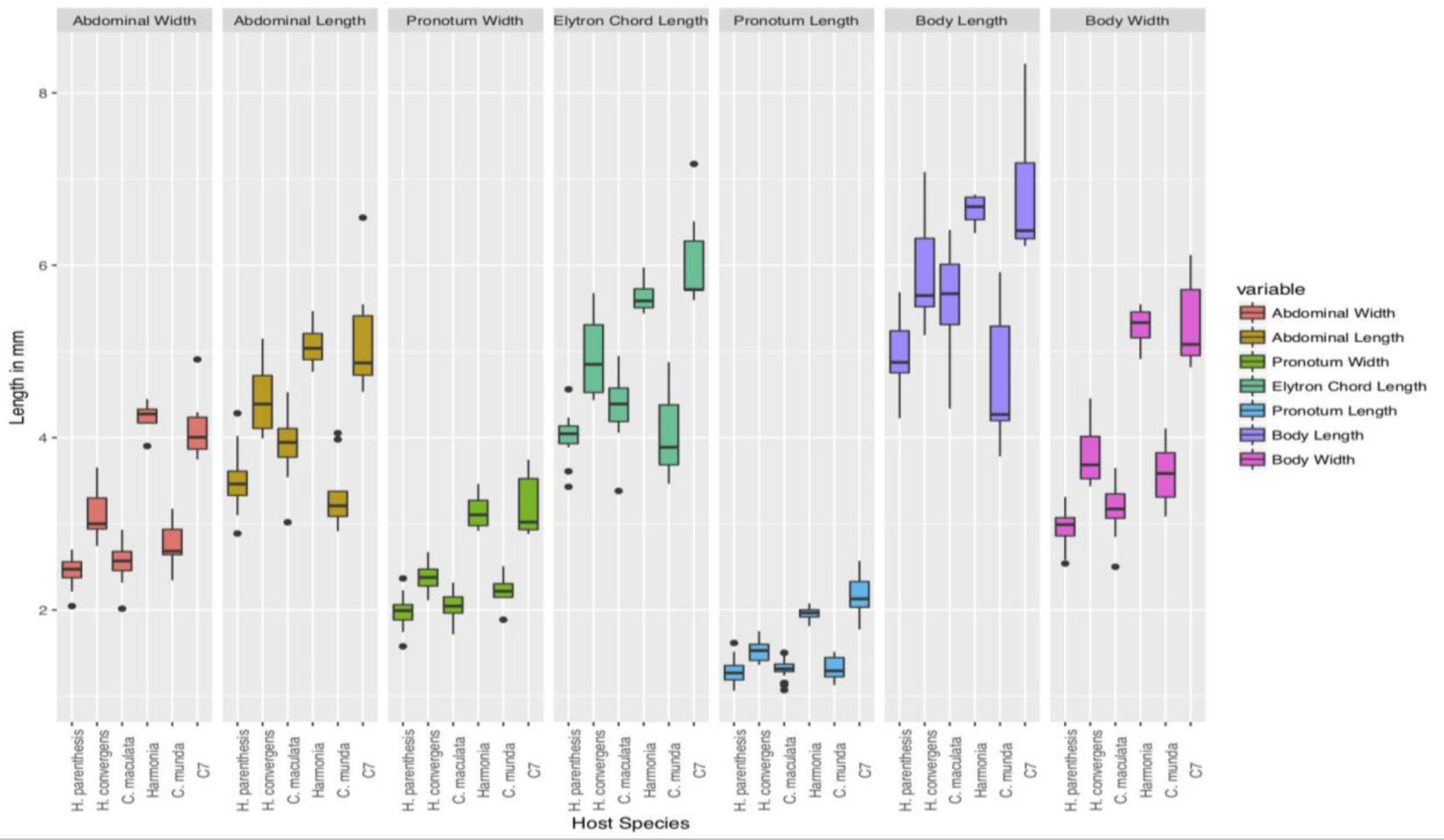
Box plot summary of Coccinellid host (*H. parenthesis, H. convergens, C. maculata, C. septempunctata, H. axyridis, C. munda*) morphometric measurements from the first round of measurements, including abdominal widths, abdominal lengths, pronotum widths, elytron chord lengths, pronotum lengths, body lengths, and body widths, in millimeters. Shown are means, and interquartile ranges within each measurement.

**Figure 5.**
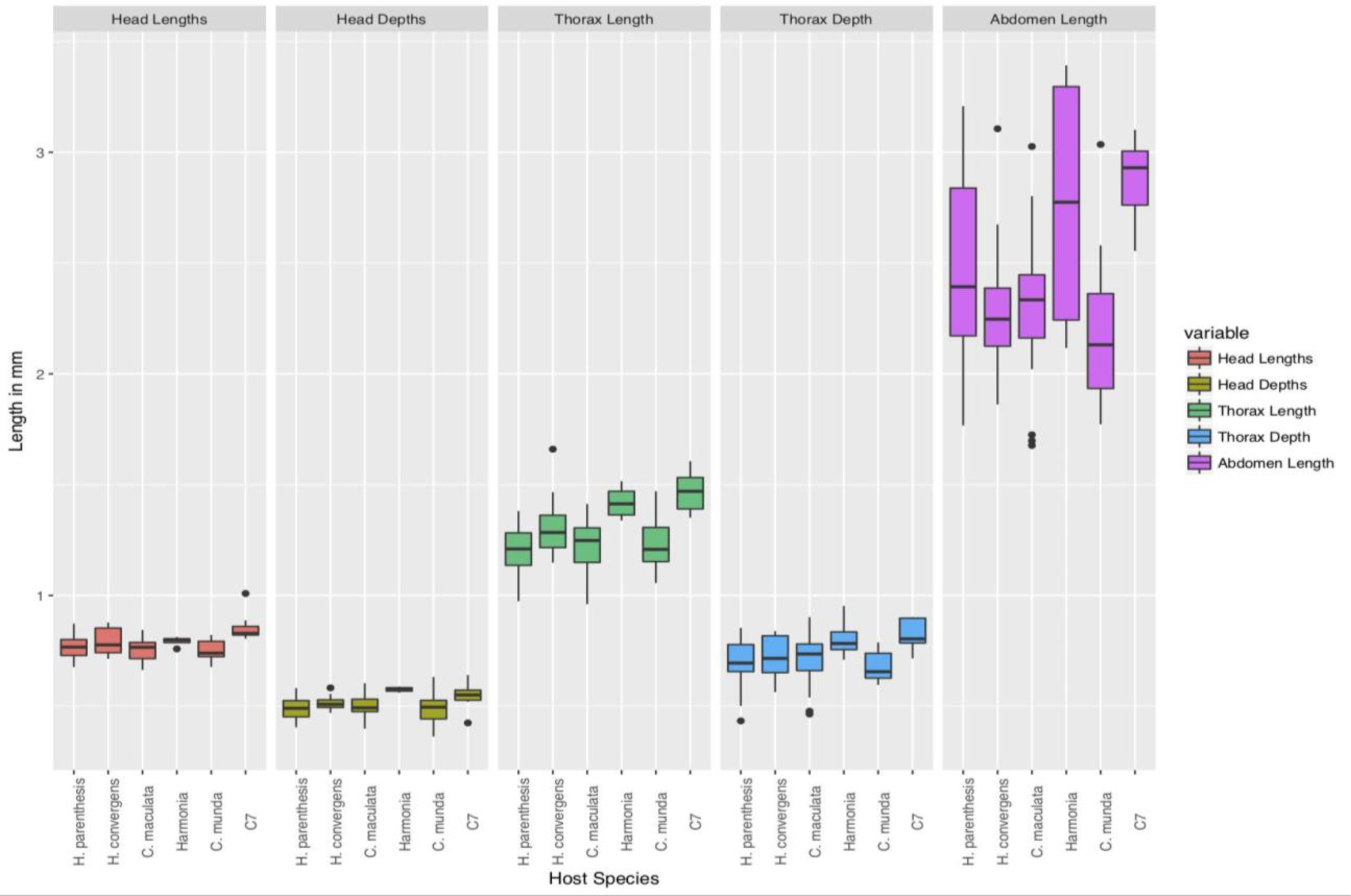
Box plot summary of parasitoid wasp (*D. coccinellae*) morphometric measurements, including head length, head depth, thorax length, thorax depth, and abdomen length, in millimeters, from the first round of measurements, categorized by their respective host species. Shown are means, interquartile ranges within each measurement. These measurements are consistent with observations on *D. coccinellae* in Table 2 of Obrycki 1988.

**Figure 6.**
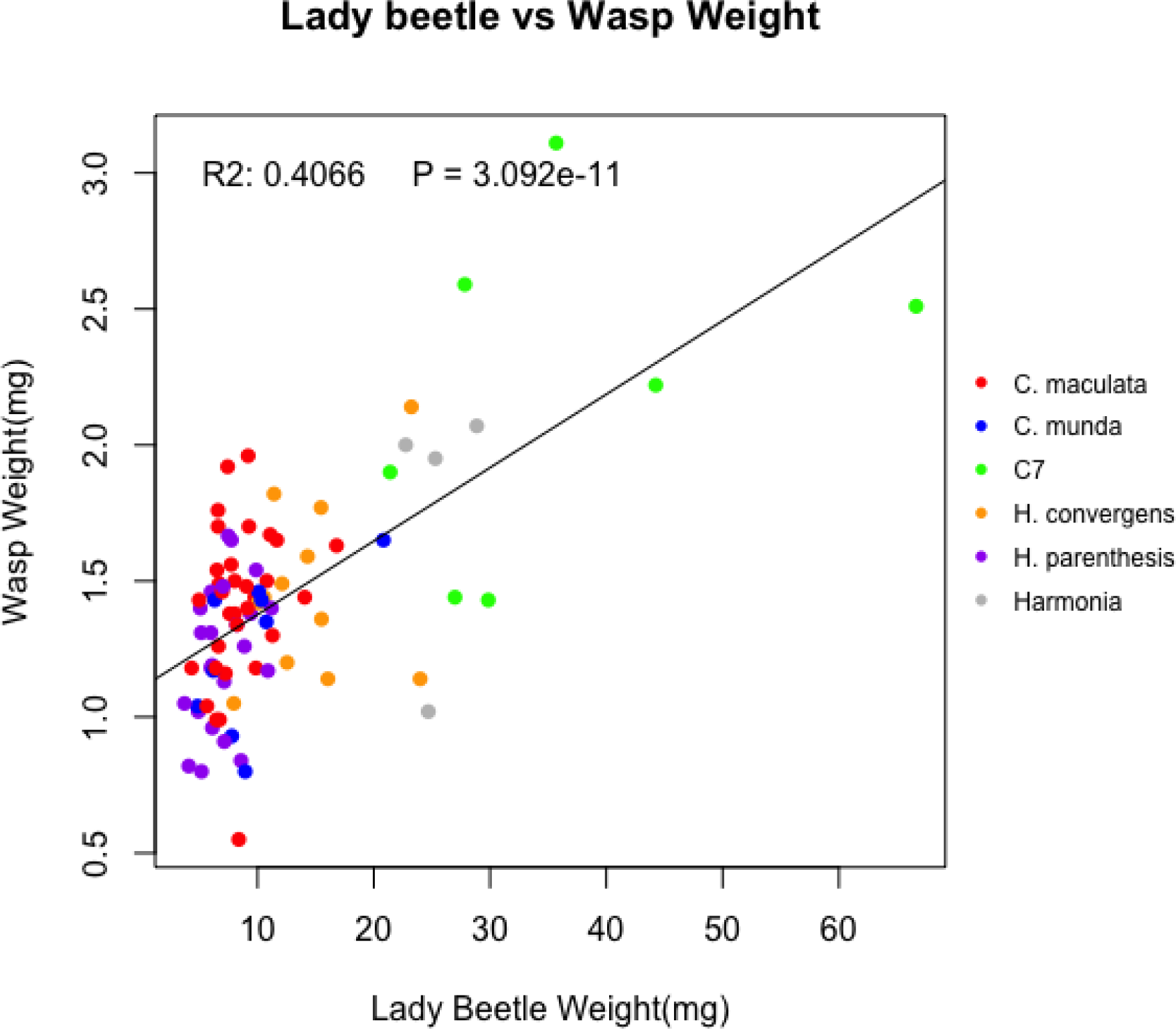
Regression of coccinellid host dry weight versus parasitoid wasp dry weight in milligrams, showing the positive correlation between size of the host and the size of its parasitoid wasp. R^2^ = 0.407, p-value = 3.09e-11.

**Figure 7.**
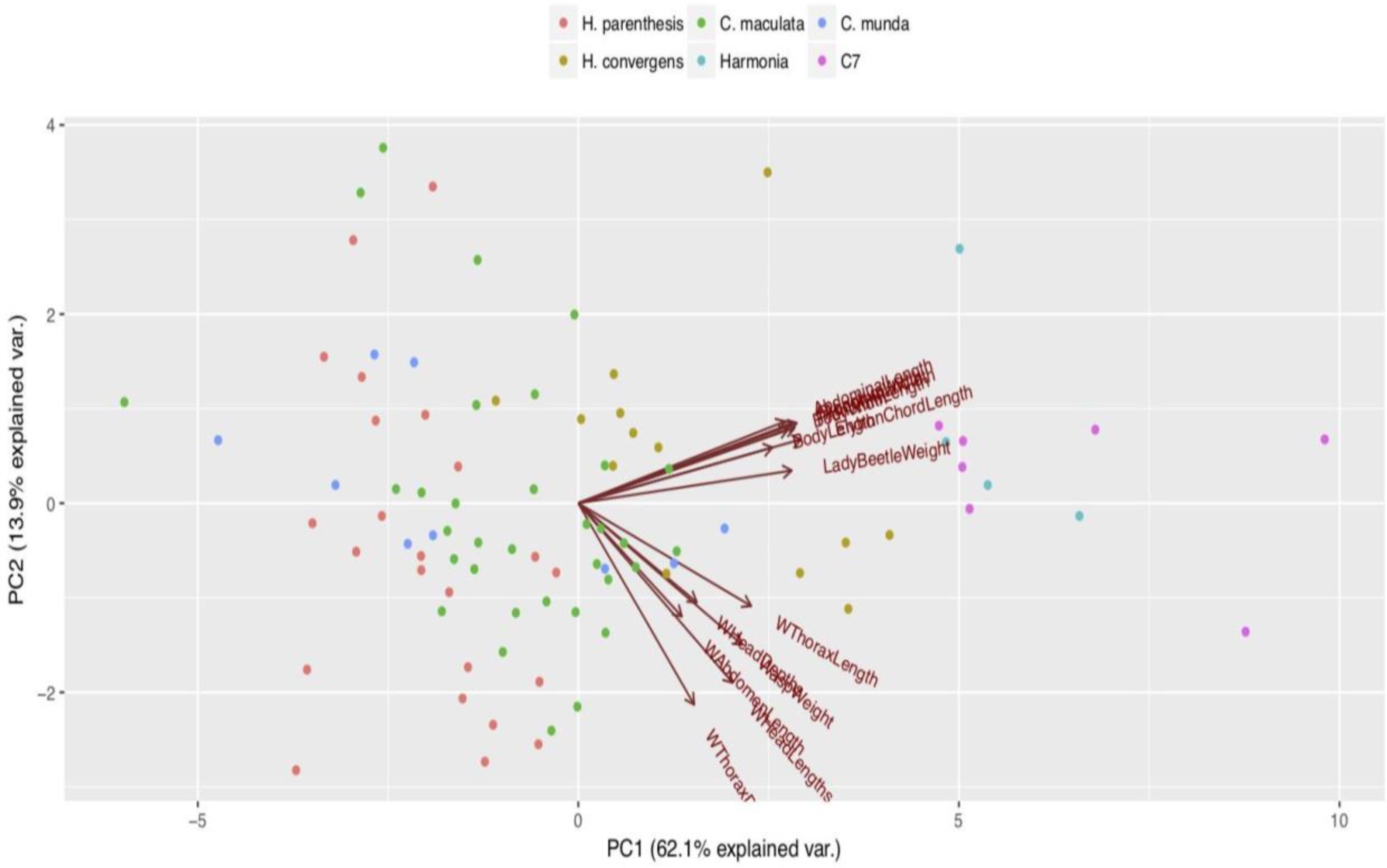
A Principal Components Analysis (PCA) plot of all variability in morphometric measurements from parasitoid wasps, and their coccinellid hosts from round 1 of measurements. The first two PC’s accounted for most of the variability in the data (PC1: 62.1%, PC2: 13.2%), with the PC1 describing variation in the coccinellid host morphometrics, and PC2 describing variation in the parasitoid wasp morphometrics.

## Discussion

The parthenogenic parasitoid *Dinocampus coccinellae* (Schrank) (Hymentoptera: Braconidae), a cosmopolitan species, attacks over 50 species in the subfamily Coccinellinae (Ceryngier et al., 2012). This parasitoid typically attacks adults, but laboratory studies and field collections of pre-imaginal stages have documented that it will attack larval and pupal stages of ladybird beetle hosts (Ware et al 2010; Obrycki et al 1985). A recent study has shown that the replication of an endosymbiotic RNA virus in the parasitoid *D. coccinellae* is correlated with the changes in host behavior following parasitization (Dheilly et al 2015). Results indicate that the manipulation of host behaviors by this parasitoid may be regulated by this endosymbiotic virus.

In our current study of the parasitoid-host interactions between *D. coccinellae* and several native North American and introduced species of Coccinellidae, we have quantified the influence of host size on the size of parasitoid females. Our study clearly shows that larger coccinellid hosts that are parasitized by *D. coccinellae* produce larger female parasitoids, an observation that has also been reported by Obrycki (1988, 1989), Belnavis (1988 - except this study indicated that larger host beetles did not always produce larger wasps). It is interesting to note that two of the larger host species examined in this study, *Cocc. septempunctata* and *Har. axyridis* are introduced species, which have established and spread throughout North America during the past four decades (Sethuraman et al 2017). It is not yet known if populations of *D. coccinellae* was introduced with *Cocc. septempunctata* or *Har. axyridis* creating a situation that may indicate that the newly introduced parasitoid populations have adaptively shifted hosts in North America since introduction in the early 20^th^ century. This possibility will be explored in a follow up study by delineating the evolutionary history and genomics of *D. coccinellae*. Nonetheless, our findings could have two potential causes - (1) morphometric variation in *D. coccinellae* is governed by phenotypic plasticity with the size of the emerging offspring contingent on the size of the coccinellid host (Boivin 2010, Benard 2004), and/or (2) morphometric variation in *D. coccinellae* is governed by genomic adaptation to its coccinellid host population (Henry et al., 2006).

Importantly, morphometric size variation in parasitoid wasps play a causal role in parasitization efficacy across their hosts. Previous studies of *D. coccinellae* parasitization efficacy in the non-native *Har. axyridis* adults have indicated the potential for behavioral adaptations in adult hosts, resulting in greater time to parasitize, when compared to the native North American species, *Col. maculata* (Firlej et al., 2009). It has also been noted that parasitization efficacy by *D. coccinellae* is significantly lower in *Har. axyridis*), when compared to conspecific native species (*Cocc. septempunctata*) in England (Comont et al., 2014), and the native species *Col. maculata* in North America (Hoogendoorn et al., 2002). However, a contradictory study of the two species that compared the parasitization rates of *D. coccinellae* of native versus introduced populations of *Cocc. septempunctata* and *Har. axyridis* in Japan and England indicated no differences based on their geographical origin, but complement the study of Firlej et al. 2009 in showing that *Har. axyridis* are parasitized at a significantly lower rate than *Cocc. septempunctata. Har. axyridis* is also known to be a voracious and invasive species across its geographical range in the world, potentially owing to behavioral and morphological adaptations to parasitization, and “enemy release” (Ceryngier et al., 2012). However, a more recent study by Dindo et al., 2016 compared the interactions of *D. coccinellae* with *Har. axyridis*, and *Adalia bipunctata* in the field, showing that *D. coccinellae* had more of a negative effect on the fitness of the *Har. axyridis* population, than on that of *A. bipunctata*. These contradicting observations in different populations of the same species indicate that behavioral, morphological, or biochemical adaptations to parasitization potentially have a genomic basis, as previously observed in *D. melanogaster* (Orr and Irving 1997). Studying the population genomic variation across coccinellid species (currently underway in the authors’ laboratories) will thus allow us to explore functional genomic variation in *Har. axyridis* and *Cocc. septempunctata* leading to defense against parasitization by *D. coccinellae*. Complementarily, we are also studying the genomics of *D. coccinellae* to study changes in the parasitoid wasp and/or intraspecific variation among populations, that allow for higher rates of successful parasitization of *Har. axyridis* in North America, or alternately, adaptively evolving to parasitize a several new host species in North America.

Body size of coccinellid hosts have also been studied to directly affect the rate of parasitization by parasitic wasps during different life history stages (*Cocc. septempunctata*, see Song et al., 2017). Since our study only controlled for the life history stage of the emerging wasp, and not for the life history stage of the coccinellid host, further studies are required to understand the efficacy of parasitization of large versus small parasitoid wasps on larval versus adult coccinellid hosts. Additionally, the sex of the coccinellid host, and prey availability in the field could also influence variability in size of adults (Belnavis 1988), which were not controlled in our study.

Our study however brings into question the fecundity of larger adult female *D. coccinellae* (presumably greater than that of smaller adult female *D. coccinellae* possibly due to more eggs and or larger eggs in larger female parasitoids and possibly longer life span of larger females, and thus greater rates of parasitization of larger hosts - see Obrycki 1989). Thus if there is indeed positive fecundity selection for larger females in a population, we would expect an ongoing trend of observing larger *D. coccinellae* in the field, which thus parasitize a larger number of native, and non-native species. Of potential interest then is the differential efficacy of parasitization of small *D. coccinellae* on larger coccinellid hosts, and vice versa. Within our 99 sampled wasps in the current study, one *D. coccinellae* female reared from a field collected *Cyc. munda* successfully parasitized and produced female F1 progeny from *Cocc. septempunctata*, *Har. axyridis* and *Col. maculata*, with the former two species being the largest of Coccinellid host species studied in this study work.

## Acknowledgments

We thank Drs. Diego Sustaita, Casey Mueller, and John Eme, Department of Biological Sciences for help with morphometric measurements. This work was funded by a CSUSM GPSM grant to the corresponding author, and by the Summer Scholars Program, CSUSM for supporting the cofirst authors, and USDA-REEU 2017-06423 Grant.

## Disclosure

The authors declare no conflict of interest, financial or other vested interests in the study species, or industrial applications of the species for biological control

## References

1 Bakker K, van Alphen JJM, van Batenburg FHD, van der Hoeven N, Nell HW et al. (1985) The function of host discrimination and superparasitization in parasitoids. Oecologia 67: 572-576. doi:10.1007/BF00790029.

2 Balduf W. V. 1926. The bionomics of Dinocampus coccinellae Schrank. Ann. Entomol. Soc. Am. 19: 465-498.

3 Belshaw R, Grafen A, Quicke DLJ (2003) Inferring life history from ovipositor morphology in parasitoid wasps using phylogenetic regression and discriminant analysis. Zool J Linn Soc 139: 213-228. doi:10.1046/j.1096-3642.2003.00078.x.

4 Benard, M. F. (2004). Predator-induced phenotypic plasticity in organisms with complex life histories. Annu. Rev. Ecol. Evol. Syst., 35, 651-673.

5 Boivin, G. (2010). Phenotypic plasticity and fitness in egg parasitoids. Neotropical Entomology, 39(4).

6 Brandl R, Vidal S (1987) Ovipositor length in parasitoids and tentiform leaf mines: adaptations in eulophids (Hymenoptera: Chalcidoidea). Biol J Linn Soc 32: 351-355. doi:10.1111/j.1095-8312.1987.tb00436.x.

7 Ceryngier, P., Nedvĕd, O., Grez, A. A., Riddick, E. W., Roy, H. E., San Martin, G.,… & Haelewaters, D. (2017). Predators and parasitoids of the harlequin ladybird, Harmonia axyridis, in its native range and invaded areas. Biological Invasions, 1-23.

8 Charnov EL, Skinner SW (1984) Evolution of host selection and clutch size in parasitoid wasps. Florida Entomol 67: 5-21. doi:10.2307/3494101.

9 Cloutier C, Duperron J, Tertuliano M, McNeil JN (2000) Host instar, body size and fitness in the koinobiotic parasitoid Aphidius nigripes. Entomol Exp Applicata 97: 29-40. doi:10.1046/j.1570-7458.2000.00713.x.

10 Cohen JE, Jonsson T, Müller CB, Godfray HC, Savage VM. Body sizes of hosts and parasitoids in individual feeding relationships. Proceedings of the National Academy of Sciences. 2005 Jan 18;102(3): 684-9.

11 Dheilly, N.M. et al 2015. Who is the puppet master? Replication of a parasitic wasp-associated virus correlates with host behavior manipulation. Proc R. Soc. B. 282:20142772. http://dx.doi.org/10.1098/rspb.2014.2772

12 Dindo, M. L., Francati, S., Lanzoni, A., di Vitantonio, C., Marchetti, E., Burgio, G., & Maini, S. (2016). Interactions between the multicolored Asian lady beetle Harmonia axyridis and the parasitoid Dinocampus coccinellae. Insects, 7(4), 67.

13 Ellers J, Van Alphen JJM, Sevenster JG (2002) A field study of sizefitness relationships in the parasitoid Asobara tabida. J Anim Ecol 67: 318-324.

14 Geoghegan, L.E., T.M. O. Majerus, & M.E.N. Majerus. 1998. Differential parasitization of adult and pre-imaginal Coccinella septempunctata (Coleoptera: Coccinellidae) by Dinocampus coccinellae (Hymenoptera: Braconidae). European J Entomology 95: 571-579.

15 Godfray, H.C.J. 1994. Parasitoids : Behavioral and Evolutionary Ecology. Princeton Univ. Press. 473 pp

16 Harvey JA, Vet LEM, Witjes LMA, Bezemer TM (2006) Remarkable similarity in body mass of a secondary hyperparasitoid Lysibia nana and its primary parasitoid Cotesia glomerata emerging from cocoons of comparable size. Arch Insect Biochem Physiol 61: 170-183. doi:10.1002/arch.20080.PubMed:16482580.

17 Hochberg, M.E. and A.R. Ives. (eds) 2000. Parasitoid Population Biology. Princeton Univ. Press. 366 pp

18 Hoogendoorn, M., & Heimpel, G. E. (2002). Indirect interactions between an introduced and a native ladybird beetle species mediated by a shared parasitoid. Biological Control, 25(3), 224-230.

19 Lee M Henry, Bernard D Roitberg, David R Gillespie Proc. R. Soc. B 2006. Covariance of phenotypically plastic traits induces an adaptive shift in host selection behaviour 273 2893-2899; DOI:10.1098/rspb.2006.3672. Published 22 November 2006

20 Mackauer, M., and A. Chau. “Adaptive Self Superparasitism in a Solitary Parasitoid Wasp: The Influence of Clutch Size on Offspring Size.” Functional Ecology, vol. 15, no. 3, 2001, pp. 335-343. JSTOR, JSTOR, www.jstor.org/stable/2656353.

21 Moore J (2002) Parasites and the behavior of animals. New York: Oxford University Press.

22 Nicol CMY, Mackauer M (1999) The scaling of host body size and mass in a host-parasitoid association: influence of host species and stage. Entomol Exp Applicata 90: 83-92. doi:10.1046/j.1570-7458.1999.00425.x.

23 Obrycki, J.J. 1989. Parasitization of native and exotic coccinellids by Dinocampus coccinellae (Hymenoptera: Braconidae). J. Kansas Entomol. Soc. 62: 211-218.

24 Obrycki, J.J., M.J. Tauber, and C.A. Tauber. 1985. Perilitus coccinellae (Hymenoptera: Braconidae): Parasitization and development in relation to host-stage attacked. Ann. Entomol. Soc. Am. 78: 852-854.

25 Orr, C.J., J.J. Obrycki, & R.V. Flanders. 1992. Host-Acceptance behavior of Dinocampus coccinellae (Shrank) (Hymenoptera: Braconidae). Ann. Entomol. Soc. Am. 85: 722-730.

26 Pennacchio, F.; Strand, M.R. Evolution of Developmental Strategies in Parasitic Hymenoptera. Annu. Rev. Entomol. 2006, 51, 233-258.

27 Sampaio, Marcus Vinicius; Bueno, Vanda Helena Paes; De Conti, Bruno Freitas. “The Effect of the Quality and Size of Host Aphid Species on the Biological Characteristics of Aphidius Colemani (Hymenoptera: Braconidae: Aphidiinae)” 2008 European Journal of Entomology, vol. 105, no. 3, p. 489.

28 Symonds MRE, Elgar MA (2013) The Evolution of Body Size, Antennal Size and Host Use in Parasitoid Wasps (Hymenoptera: Chalcidoidea): A Phylogenetic Comparative Analysis. PLoS ONE 8(10): e78297. https://doi.org/10.1371/journal.pone.0078297

29 Traynor RE, Mayhew PJ (2005) A comparative study of body size and clutch size across the parasitoid Hymenoptera. Oikos 109: 305-316. doi:10.1111/j.0030-1299.2005.13666.x.

30 Ware, R., L-J. Michie, T. Otani, E. Rhule, & R. Hall. 2010. Adaptation of native parasitoids to a novel host: the invasive coccinellid Harmonia axyridis. IOBC/WPRS Bull. 58: 175-182.

